# Allele Specific Expression in Human – Genomic Makeup and Phenotypic Implications

**DOI:** 10.1101/757997

**Authors:** Kerem Wainer-Katsir, Michal Linial

## Abstract

The allele-specific expression phenomenon refers to unbalanced expression from the two parental alleles in a tissue of a diploid organism. AlleleDB is a high-quality resource that reports on about 30,000 ASE variants (ASE-V) from hundreds of human samples. In this study, we present the genomic characteristics and phenotypic implications of ASE. We identified tens of segments with extreme density of ASE-V, many of them are located at the major histocompatibility complex (MHC) locus. Notably, at a resolution of 100 nucleotides, the likelihood of ASE-V increases with the density of polymorphic sites. Another dominant trend of ASE is a strong bias of the expression to the major allele. This observation relies on the known allele frequencies in the healthy human population. Overlap of ASE-V and GWAS associations was calculated for 48 phenotypes from the UK-Biobank. ASE-V were significantly associated with a risk for inflammation (e.g. asthma), autoimmunity (e.g., rheumatoid arthritis, multiple sclerosis, and type 1 diabetes) and several blood cell traits (e.g., red cell distribution width). At the level of the ASE-genes, we seek association with all traits and conditions reported in the GWAS catalog. The statistical significance of ASE-genes to GWAS catalog reveals association with the susceptibility to virus infection, autoimmunity, inflammation, allergies, blood cancer and more. We postulate that ASE determines phenotype diversity between individuals and the risk for a variety of immune-related conditions.

## Introduction

In humans as a diploid organism, each individual carries two alleles for each genomic variant. Both alleles (homozygous or heterozygous), contribute to the individual’s phenotype. However, an imbalanced expression of the two alleles negates the equal contribution of the alleles to the phenotype. Allelic Specific Expression (ASE) is considered when the expressed genes exhibit unbalanced expression from the two alleles with one allele expresses in access relative to the other (Panousis et al. 2014; Reinius and Sandberg 2015).

The ASE phenomenon is a special case of expression quantitative trait loci (eQTL). eQTL are regulatory SNVs that affect the amount of expression from a gene. They are detected by the variability of expression across many tissues and individuals (Gilad et al. 2008, Consortium, 2017 #3). ASE is best explained by cis-regulation of eQTL which stand for a genomic variant that is positioned at the same chromosome as the target gene, affecting its expression (Hasin-Brumshtein et al. 2014; Hu et al. 2015). Most likely, a polymorphic site at the vicinity of the gene alters the binding affinity or specificity of a transcription factor that regulates its expression (Boyle et al. 2012; Karczewski et al. 2013, Chen, 2016 #1).

The most known permanent ASE phenomena include Chromosome X-inactivation (Pollex and Heard 2012; Lee and Bartolomei 2013) and imprinted genes (DeVeale et al. 2012; Babak et al. 2015; Baran et al. 2015b). In the former, expression from the inactivated chromosome is suppressed as a result of a non-reversible change in chromatin structure (Avner and Heard 2001; Galupa et al. 2015). Epigenetic signature governs the parent-dependent expression for imprinted genes (Babak et al. 2015; Chuang et al. 2017; Santoni et al. 2017).

Technically, determining ASE variants and genes is based on quantifying the expression levels from heterologous single nucleotide variants (hSNVs) (Pandey et al. 2013), using alignment of RNA-seq data to the genome. This procedure is sensitive to data quality, sequencing depth (Skelly et al. 2011; Conesa et al. 2016) and additional biases (discussed in (Hodgkinson et al. 2016)). For example, mapping of transcribed sequenced reads to the reference genome led to an inherent bias toward the reference allele, and underrepresenting of the alternative allele (Degner et al. 2009; Stevenson et al. 2013). Furthermore, determining genes as ASE in single cells (Zhang et al. 2009; Rozowsky et al. 2011) is prone to a large technical noise due to the stochastic nature of expression from single cells. This stochastic nature is largely explained by the phenomenon of transcriptional bursting that leads to monoallelic expression (MAE) (Larsson et al. 2019). Moreover, transcripts’ dropout (Islam et al. 2014) tend to mask the ASE signal in single cells (Kim et al. 2015).

Measuring tissue expression given the actual polymorphic map of individual genomes is a valuable resource to study the extent of ASE is each tissue. The estimates for the ASE phenomenon span over a range of organisms (Chamberlain et al. 2015; Crowley et al. 2015). The human species genomics research is unique in having the largest collection of sequenced variants (Chen et al. 2016). GTEx database studied ASE in 44 sequenced tissues and across 175 individuals (Consortium 2015). Compilation of data across many individuals in GTEx reveals that ∼50% of all genes (with a relaxed definition of ±1M bp) show evidence for cis-regulation, with only a small fraction of them being genuine ASE (1.7%-to 3.7%), with the highest fraction in whole blood samples (Consortium 2015). From a catalog of genes from 13 types of immune cells, 41% show cell-type-specific expression that is linked to cis-associated variants (Schmiedel et al. 2018), and ∼5% show it in human primary T-cells (Heap et al. 2010). ASE-V were associated with various human phenotypes, as shown for numerous neurodevelopmental diseases (e.g. (Jeffries et al. 2013; Huang et al. 2017)).

Studying the genetic basis of complex diseases and phenotypes is based on seeking statistical differences of case-control cohorts (Sudlow et al. 2015). The results of such genome-wide association studies (GWAS) are SNVs which are statistically associated with the studied phenotypes (MacArthur et al. 2016; Bycroft et al. 2017). The majority of GWAS associations are attributed to cis-regulatory variants (Bonder et al. 2017; Consortium et al. 2017; Barbeira et al. 2018) that affect the expression of coding and non-coding genes (Manke et al. 2010; Maurano et al. 2015). It is anticipated that a fraction of these variations in the population exhibits ASE and therefore impact phenotypic diversity (De La Chapelle 2009; Heap et al. 2010).

A high-quality, exhaustive resource for human ASE variants and genes is AlleleDB (Chen et al. 2016). It is based on RNA-seq and Chip-seq samples from lymphoblastoid cell lines (LCL) form 382 individuals with complete genomes (Chen et al. 2016). In this report, we used AllelDB for characterizing the ASE phenomenon given the genome organization and show the association of ASE-V with a range of human phenotypes. We have characterized the increase likelihood of ASE to occur at highly polymorphic clusters across the genome, and the tendency of ASE to the major allele. We observed the majority of ASE variants to be associated with determinants of the immune system. Based on GWAS association studies for human traits and diseases we illustrate that ASE genes are associated with an enhanced risk for autoimmunity, inflammatory diseases, and viral infection.

## Results

### ASE variants are widely spread on all autosomal chromosomes

We set to characterize the appearance of ASE variants (ASE-V) across the human genome. To this end, we analyze data from a high-quality resource of ASE-V from human LCL (AlleleDB, (Chen et al. 2016). Out of all accessible expressed hSNVs (abbreviated Acc-V), those with strong statistical evidence for an unbalanced allelic expression are labeled allelic specific expressing SNVs (abbreviated ASE-V) (Chen et al. 2016). The final collection of the unique set of SNVs includes 287,453 Acc-V and 29,346 ASE-V.

Fig. 1 is a Manhattan-like plot of the 22 autosomal chromosomes for the non-redundant ASE-V collection. The value associated with each ASE-V (supported by >1 individual, total 29,346) is the -log(p-value) of the beta-binomial significance of being ASE. Notably, ASE sites are distributed over all autosomal chromosomes. However, some chromosomal loci are especially dense with significant ASE-V.

**Figure 1.**
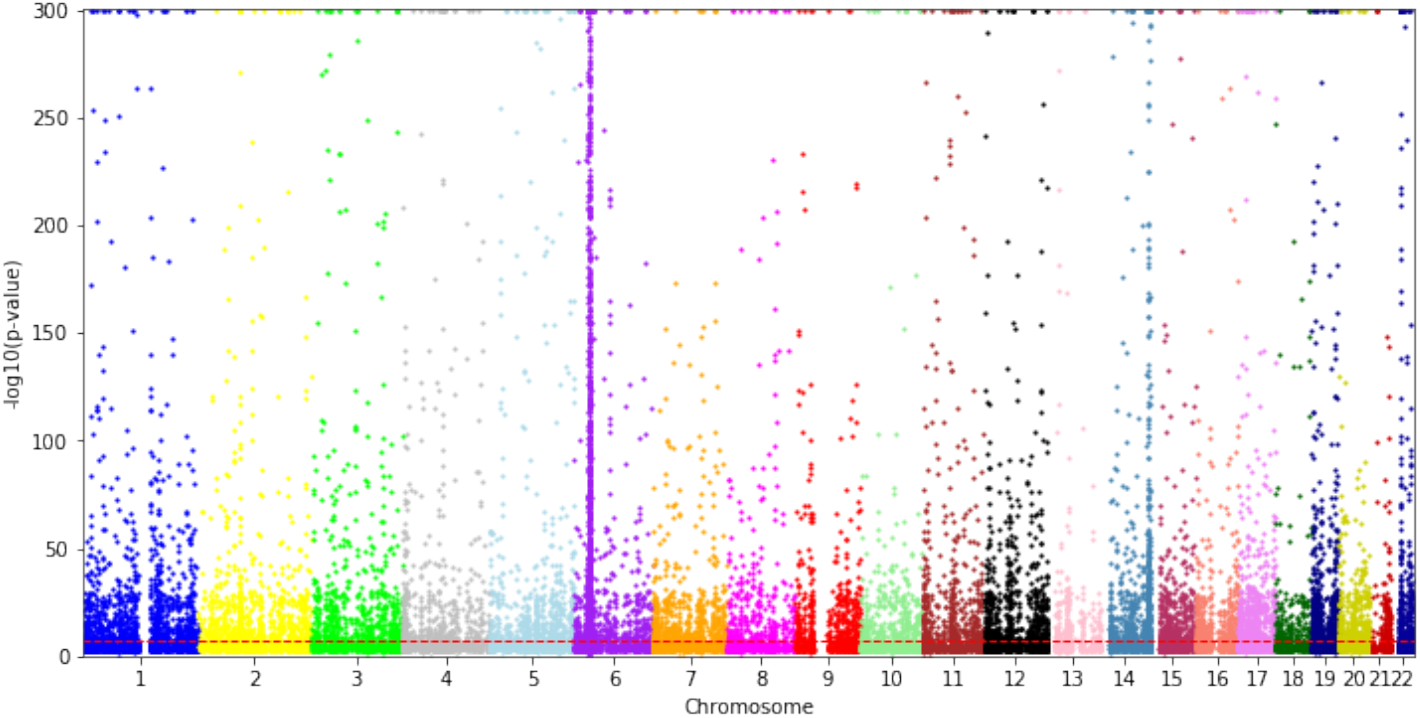
A Manhattan-like plot of ASE-V. The genomic coordinates are displayed and colored for the autosomal chromosomes (excluding ChrX, ChrY and mitochondria chromosome). The y-axis is the -log (p-value) of the beta-binomial significance of the selected SNVs. The plot reports 29,346 SNVs (based on AlleleDB, (Chen et al. 2016)). Only SNVs with minimal support of >=2 individuals are included. The confidence for some of the variants is extremely high with 473 ASE-V having p-value < 1.0E-300. Genomic regions with dense representations of ASE-V occur at numerous chromosomes (e.g., the p-arm of Chr6).

### ASE-rich segments are not equality distributed along chromosomes

Next, we tested the possibility that the appearance of ASE-V reflects a simple sampling of Acc-V. Fig. 2A provides a view at a chromosomal resolution of 100k bp segments for the ratio of ASE-V to Acc-V for three representative chromosomes. Certain chromosomal regions are particularly dense with high ASE-V to Acc-V ratio (e.g., the p-arm of Chr6, the tail of Chr14 q-arm, Fig. 2A). Supplementary Figure S1 expands the view for all autosomal chromosomes. To highlight the chromosomal positional properties in term of ASE occurrences, we created a cumulative profile of sequential 100k bp segments for each chromosome. The plots in Fig. 2B capture the rate of accumulation of segments with Acc-V (x-axis) and ASE-V (y-axis). Chromosomes in which the accumulation of ASE and Acc segments occur at a similar pace imply unbiased sampling (e.g., Chr1). In contrast, chromosomes that show a strong irregularity of the scatter plot are consistent with over-representation regions with ASE-segments. Supplementary Figure S2 shows the scatter plots for the cumulative view for ASE-V and Acc-V for all chromosomes.

**Figure 2.**
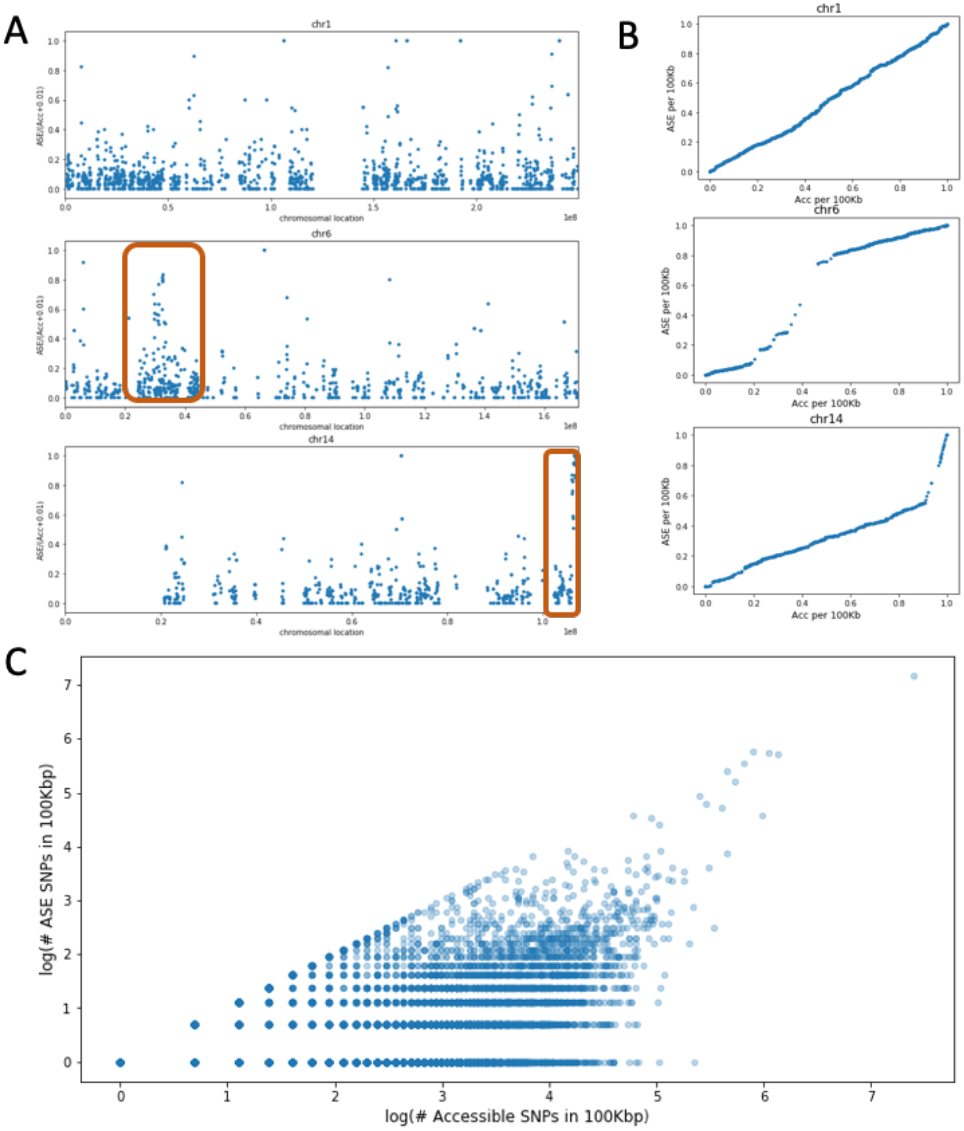
Genomic representation of ASE-V rich segments. (A) Chromosome representation with the ratio of ASE-V to Acc-V in segments of 100k bp. For visualization clarity, only instances with ≥10 Acc-V per 100k bp segment are included. A local concentration of segments with a high ratio is indicative of a rich chromosomal region that extends the 100k segment scale. Few of such regions are highlighted (red frames). (B) A cumulative view of 3 representative chromosomes. The axes show the cumulative fraction of segments with Acc and ASE, respectively. Segments with no indication of expressed SNVs are compressed. The irregularity in the scatter plot is indicative of a locally concentrated of ASE rich segments. For example, in Chr14, 45% of the ASE segments (y-axis, 0.55 to 1.0) match only 10% of the Acc segments. (C) A scatter plot for all 17,100 genomic segments (100k bp each) with evidence for expression (i.e. there is at least one evidence for Acc-V in a segment). These segments are presented in a log-scale. Note the extreme instances (at the top right corner) with a large number of Acc-V and strong evidence of ASE-V.

A statistical Kolmogorov–Smirnov (KS) test was applied to compare the distribution probability for each chromosome (Supplemental Table S1). The significant results of the KS test for all chromosomes allow rejecting the hypothesis that ASE-V is a simple sampling of the Acc-V. Notably, maximal confidence for this test is associated with Chr6, Chr9, and Ch14. The statistical significance of Chr9 is due to a long segment which is ASE-V depleted.

The overall statistics of the ASE-rich genomic segments at a resolution of 100k bp is shown in Fig. 2C. All autosomal chromosomes were analyzed through 100k bp segments (with a 50k bp overlapping windows). Altogether, there are 17,100 segments with Acc-V evidence which account for 29.7% of all tested 100k bp segments. On average, the number of Acc-V per 100k bp segment is 16.8. Fig. 2C shows a scatter plot of all segments according to the number of Acc-V and ASE-V (log-scale). Half (51%) of these segments have <10 Acc-V in a segment. Considering only segments with dense Acc-V (≥100) highlights two extreme observations, segments with no evidence for ASE-V that will not be further discussed, and segments that are ASE-V-rich.

### Genomic loci rich with ASE segments are associated with immune gene clusters

The appearance of Acc-V and ASE-V in a 100k bp segment highlighted several extreme dense segments (Fig. 2C, top right). The richest segment of all genome has 1645 Acc-V and 1313 ASE-V. This segment belongs to Chr6 within the major histocompatibility complex (MHC) and covers the HLA-DRB1 and HLA-DQA1 genes. These gene products belong to MHC-Class II that act to display foreign peptides to the immune system. Altogether we report on 166 instances in which the number of Acc-V is ≥100 (Supplemental Table S2). Table 1 lists the properties of the 30 most dense ASE-V segments (with >20% of the Acc-V being ASE-V). A large fraction (43%) of these segments derives from MHC locus. Interestingly, many of the other segments that are external to the MHC locus are rich in gene clusters of various cell receptors, antibodies and immune-cell recognition related functions. For example, the BTN3A2 and BTN2A2 belong to a collection of B7/butyrophilin-like group which is a subset of the immunoglobulin (IG) gene superfamily. The cluster of BTN3 and BTN2 is considered an extension to the MHC locus (Rhodes et al. 2001). Members of this gene cluster inhibit T cell proliferation and T-cell receptor (TCR) activation, thus play a role in the adaptive immune system. Another ASE-rich segment is positioned at Chr14 (positions 106.1M-106.2M) which overlaps with an elaborate gene collection of IG heavy chain type (i.e. IGHC, IGHE, IGHG). A segment in Chr2 highlights a cluster of genes and segments for creating the light chain of IG kappa (IGKV, IGKJ). Other genes on Chr19 are members of a large cluster of leukocyte IG-like receptors (LILR, also called CD85) that are predominantly expressed in both myeloid and lymphoid lineage. These large family of receptors exerts immunomodulatory effects on a wide range of cells of the innate and adaptive system. MHC Class I recognition by LIRL proteins prevents the immune system from self-attack and as such, LIRL gene expression plays a key role in pregnancy, transplantation, arthritis and more (Hirayasu and Arase 2015). We conclude that the majority of the dense segments of ASE-V are associated with gene families acting at multiple domains of cell recognition and specificity of the immune system (via antibodies, receptors, and antigens).

**Table 1.** Genomic segments of 100k bp that are rich with ASE-V

| Chr | position<br>(in M)<sup>a</sup> | ASE in<br>100k | Acc in<br>100k | Ratio<br>ASE/Acc | Rep.<br>Gene<sup>b</sup> | Immune function |
| --- | --- | --- | --- | --- | --- | --- |
| chr2 | 89.15 | 83 | 152 | 0.55 | IGKV | IG kappa light chain |
| chr3 | 122.40 | 23 | 115 | 0.20 | PARP14 |  |
| chr4 | 5.80 | 92 | 142 | 0.65 | EVC |  |
| chr5 | 0.45 | 36 | 133 | 0.27 | AHRR |  |
| chr5 | 96.10 | 34 | 137 | 0.25 | ERAP1 | Processing HLA class 1 |
| chr6 | 26.35 | 46 | 135 | 0.34 | BTN3A2 | IG superfamily |
| chr6 | 29.85 | 112 | 273 | 0.41 | HLA-G | HLA Class I |
| chr6 | 29.90 | 308 | 463 | 0.67 | HLA-A | HLA Class I |
| chr6 | 31.10 | 51 | 155 | 0.33 | TCF19 | HLA Class I |
| chr6 | 31.20 | 221 | 287 | 0.77 | HLA-C | HLA Class I |
| chr6 | 31.30 | 141 | 222 | 0.64 | HLA-B | HLA Class I |
| chr6 | 31.35 | 25 | 113 | 0.22 | MICA | HLA Class I |
| chr6 | 32.45 | 254 | 335 | 0.76 | HLA-DRA | HLA Class II |
| chr6 | 32.50 | 318 | 367 | 0.87 | HLA-DRB5 | HLA Class II |
| chr6 | 32.55 | 312 | 425 | 0.73 | HLA-DRB1 | HLA Class II |
| chr6 | 32.60 | 1313 | 1645 | 0.80 | HLA-DQA1 | HLA Class II |
| chr6 | 32.75 | 37 | 171 | 0.22 | HLA-DQB2 | HLA Class II |
| chr6 | 33.00 | 97 | 400 | 0.24 | HLA-DOA | HLA Class II |
| chr6 | 33.05 | 121 | 237 | 0.51 | HLA-DPA1 | HLA Class II |
| chr7 | 1.05 | 32 | 151 | 0.21 | COX19 |  |
| chr9 | 37.85 | 24 | 100 | 0.24 | DCAF10 |  |
| chr14 | 24.90 | 33 | 123 | 0.27 | CBLN3 |  |
| chr14 | 106.20 | 98 | 119 | 0.82 | IGHG1 | IG gamma heavy chain |
| chr14 | 106.30 | 181 | 307 | 0.59 | IGHD | IG delta heavy chain |
| chr16 | 2.00 | 23 | 113 | 0.20 | RPL3L |  |
| chr17 | 3.55 | 43 | 113 | 0.38 | CTNS |  |
| chr19 | 50.40 | 35 | 113 | 0.31 | IL4I1 |  |
| chr19 | 55.10 | 38 | 137 | 0.28 | LILRA1 | Leukocyte IG receptor |
| chr19 | 58.35 | 31 | 147 | 0.21 | ZNF419 |  |
| chr22 | 44.55 | 24 | 100 | 0.24 | PARVG |  |

<sup>a</sup>The position refers to the center of the 100k bp segment. A representative gene is a selected
gene from the discussed segment.

In addition, several other highly polymorphic ASE-rich segments (Table 1) show no direct link to any exceptional immune-related loci or gene clusters. Two of the most dominant instances associate with Chr4 (position 5.8M) and Chr17 (position 3.55M) (Table 1). Supplemental Fig. S3A shows the position of ASE-V on Chr4 segment (5.75M-5.85 M). Two genes are included in this segment EVC and CRMP1. The ASE signal is localized mostly at the 3’-UTR of the EVC. A peak of expression at this location was also documented by NK cells, T cells-CD4 and classical monocytes as reported by the DICE database (Schmiedel et al. 2018). We attribute the extreme signal of the EVC gene to its characteristics as a non-classical imprinted gene. Specifically, EVC belongs to a small set of genes that exhibit a monoallelic expression (MAE) that is not deterministic among individuals (Baran et al. 2015a). A similar observation is associated with P2RX5 on Chr17 (Supplemental Fig. S3B). The100k bp segment of Chr17 (3.5M-3.6M) is gene rich and the ASE-V overlaps P2RX5 (and P2RX5-TAX1BP3 readthrough) that is expressed on B-cell and activated T-cells (Schmiedel et al. 2018). P2RX5 was specified by its random MAE (Gimelbrant et al. 2007). We conclude that high-density of ASE-V in genomic segments covers the molecular building blocks of the immune system including antigen presentation, immunoglobulin (IG), TCR and Leukocyte receptors, in addition to instances of permanent MAE genes.

### Accessible variants cluster in dense polymorphic short segments

We seek to quantify the genomic characteristics of Acc-V occurrences. To this end, we measured the spacing among Acc-V in the genome. Fig. 3A shows the fraction of the minimal spacing of Acc-V with respect to a theoretical geometrical model (see Methods). The bar plot in Fig. 3A shows how many of the Acc-V spacing falls within a predefined segment length for Chr1 (as a representative). The preference of Acc-V to appear in a cluster in surprisingly high and dominated by short-range distances of 50 and 100 bp. A trend of Acc clustering within short distances applies for any distances up to 1000 bp. The deviation from expectation is maximal for short spacing (Fig. 3B) with 12.5 folds and 9.75 folds difference for spacing up to 50 and 100 bp, respectively. Consequently, segments of very long distance spacing (>20,000 bp) are less abundant relative to the calculated expectation. All reported distances (5835 distances >=1 bp, Chr1) are listed in Supplemental Table S3. We show that Acc-V tend to cluster in very compact local sequences. These define the prevalence of Acc-V to highly polymorphic regions in the genome.

**Figure 3.**
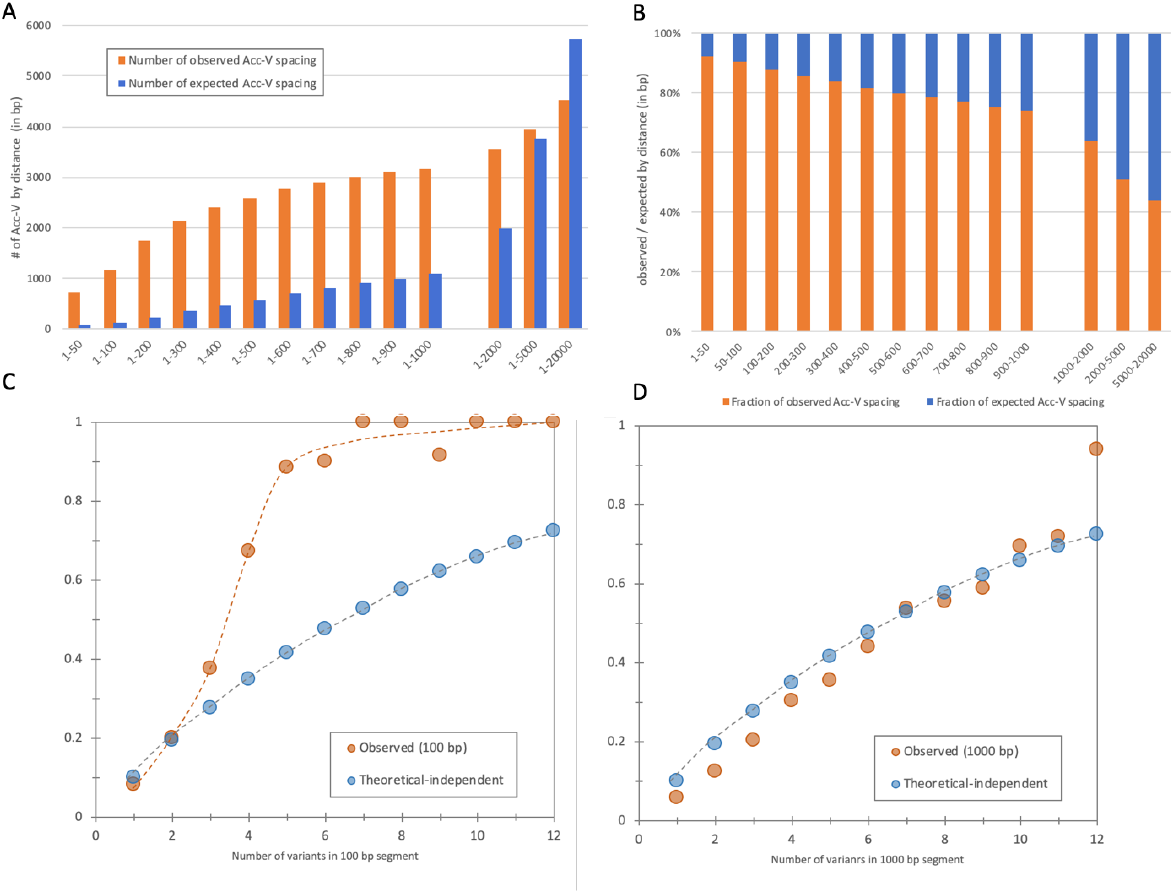
Characteristics of clustering and spacing tendency for ACC-V and ASE-V. (A) The bar plot shows results for Chr1 for the minimal distances of Acc-V to a consecutive one. The distances tested ranges from 1-50 bp to 1-20,000 bp. The expectation distances are based on the geometric expectation for Chr1 according to the global density of Acc-V in the chromosome (see Methods). (B) The relative fraction of observed Acc-Vs out of the observed and expected in each range of the spacing distances. For example, for a 1-50 bp spacing distance, enrichment of the observed versus the expected is maximal (12.1 folds). The analysis is based on 5835 reported distances (>=1 bp, Chr1) listed in Supplemental Table S3. (C) The likelihood of a site to include ASE-V given the number of polymorphic sites (Acc-V) recorded for a length of 100 bp. (D) The likelihood of a site to include ASE-V in view of the number of polymorphic sites (Acc-V) recorded for the length of 1000 bp. The expected occurrence is calculated according to a probabilistic independent model (see Methods).

### The local abundance of polymorphic sites increases the likelihood for ASE

To understand the features that specify ASE occurrences, we tested the tendency of a variant to be assigned as ASE as a result of the local density of Acc-V. Based on the observation of the local clustering of Acc-V at short distances (Figs. 3A, 3B), we tested the likelihood of a site to be ASE given the number of polymorphic sites recorded (within a 100 bp window). In contrast to the arithmetic calculated expectation, the observed ASE-V show a strong cooperative trend with respect to the number of ASE-V within a local density of Acc-V (Fig. 3C). The differences are maximal at the higher densities of Acc-V. For example, having 5 Acc-V in a 100 bp segment is expected to associate with an ASE fraction of 40%, but, it accounts for a fraction of 90% (Fig. 3C). However, the signal fades at a longer distance (1000 bp, Fig. 3D). This observation supports the notion that ASE is stabilized in short segments which are prone to extensive sequence diversity at the level of individuals and population.

### ASE sites are primarily associated with the major allele

It is expected that even for the phenomenon of ASE in which the expression from an individual is validated to be strongly uneven, the overall distribution of the selected allele across the population is expected to be balanced (with the exception of imprinted genes in which parent of origin is predetermined). To test this hypothesis, we assigned each representative expressed hSNV that is labeled ASE-V to the Reference (Ref) or the Alternative (Alt) allele for each individual, according to the majority of its expressed reads being from the Ref or Alt allele. Additionally, we assign each SNV to its known population alternative allele frequency (AF, see Methods). Fig. 4A shows that while the total number of Acc-V are almost equality spread between the two alleles for each of the AF bins, the occurrences are biased for the subset of ASE-V. The calculated coefficient by a generalized linear model (GLM) for the Acc-V is 0.26, while it is much higher for ASE-V (3.11, p-value <=E-300). Therefore, the frequency of expressing from the Ref allele in ASE-V is strongly associated with the rarity of the allele. ASE-V bins by their allele frequency are listed in Supplemental Table S4.

**Figure 4.**
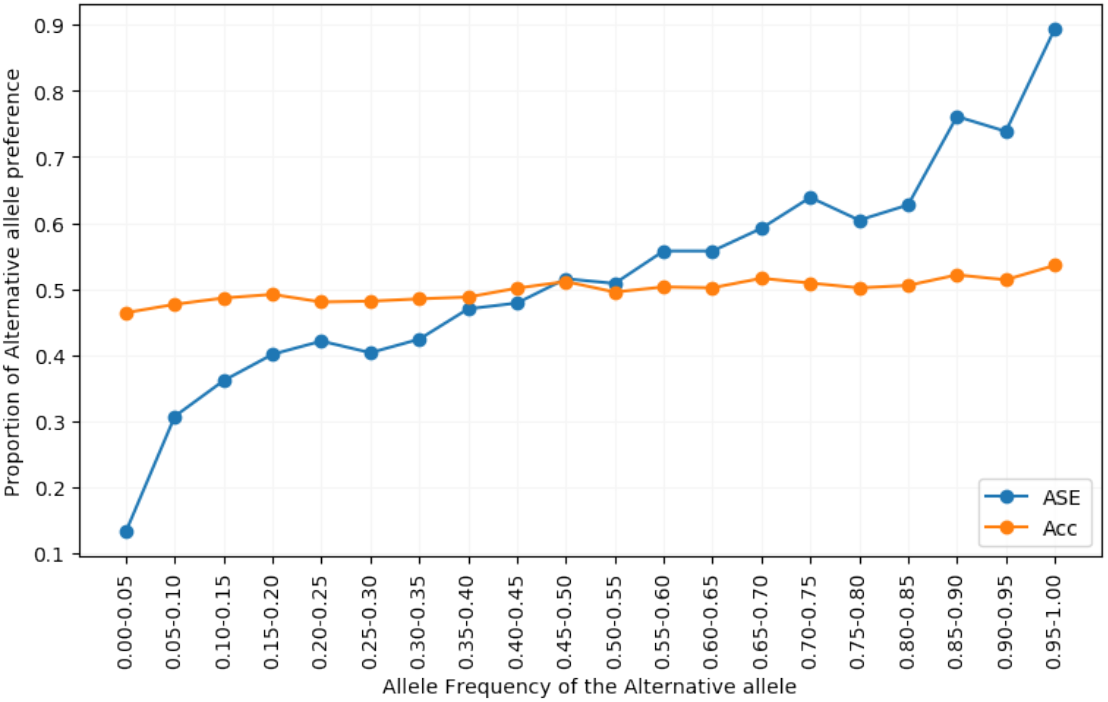
The proportion of Alternative allele expression according to Allele Frequency (AF) of the Alternative allele. ASE-V (blue) and Acc-V (orange) were binned according to their Alternative Allele Frequency (x-axis). The proportion of SNVs preferably expressed from the Alternative allele is shown on the y-axis.

We imply that the imbalance in the expression of genes associated with ASE-V is not only associated with hypermutated local regions in the genome, but it is also biased towards the expression from the major allele in the population.

### ASE sites are associated with autoimmune diseases (AID) and blood cell related traits

To test the relevance of the ASE enriched chromosomal regions to human phenotypes, we tested the association of ASE-V to a set of phenotypes and diseases. We performed a GWAS that covers 332,709 samples from the UK Biobank (UKBB) (Sudlow et al. 2015) for continuous (e.g. body mass index) and binary (e.g., diagnosed a Type 1 diabetes) phenotypes (see Methods). Altogether, we provide sets of associated variants for 48 phenotypes (Supplemental Table S5).

For each phenotype, the association of SNVs to the phenotype was tested. The significance of the SNVs association was then associated with ASE-V using Fisher Exact test (see Methods). The OR indicates the effect of the association between the significant ASE-V and the tested association (for each of the phenotype / disease). A summary of the OR results and the 95% interval confidence for significant associations (p-value of <0.05) is shown in Fig. 5A. All association results are summarized in Supplemental Table S5_GWAS.

**Figure 5.**
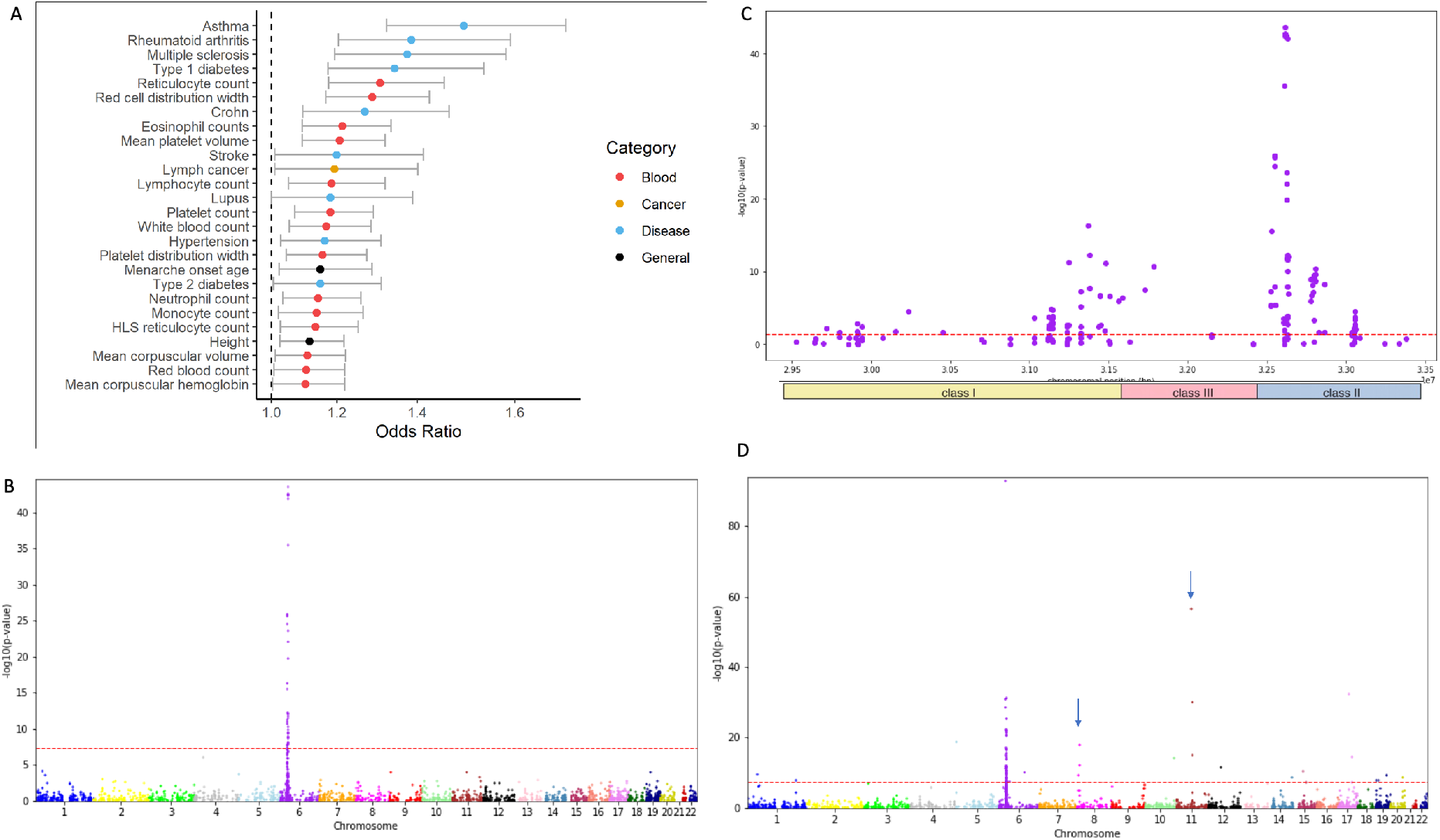
Phenotype association with ASE-V based on GWAS results for 48 major phenotypes from the UKBiobank. **(A)** A list of 25 significantly enriched by ASE-V phenotypes (p-value<0.05) ranked according to their odds ratio (OR) of SNVs being both ASE and associated with the phenotype. The 95% interval confidence for each phenotype is shown. The phenotypes are color coded for autoimmune and inflammatory diseases (denoted as Immune), Physical (e.g. height), Blood (e.g. reticulocyte count), Cancer, and Others (including type 2 diabetes, hypertension and stroke). **(B)** Manhattan plot of the association of ASE-V with asthma (OR=1.50, p-value=1.27E-10) are shown for their position on the autosomal chromosomes. **(C)** Zoom-in on the signal of asthma shown in (B) for Chr6 ∼4M bp from 29.5M to 33.5M of the MHC locus. The partition of this locus to class I, class III and class II of the MHC is shown. **(D)** Manhattan plot of the association with ‘Red cell distribution width of ASE-Vs is shown according to SNV positions on the autosomal chromosomes. The arrows mark several clustered ASE-V.

The most significant signal for the ASE-V is associated with a diverse collection of immune-associated conditions, mostly inflammatory and autoimmune diseases (AID) including asthma, type 1 diabetes (T1D), multiple sclerosis, inflammatory bowel disease (IBD) / Crohn’s disease. An additional group of phenotypes that are highly enriched includes features of blood cells. These phenotypes include the number of erythrocytes, granulocytes (e.g. eosinophils, neutrophils) lymphocytes, thrombocytes and numerous quantitative properties (width, counts, volume) of these cells.

From all six major cancer types in the UKBB population (including prostate, breast, and lung cancer (Supplemental Table S5), only blood lymphoma is significantly associated with ASE-V. Mental conditions and diseases (schizophrenia, depression) or metabolic conditions (BMI) were insignificant according to this test (Supplemental Table S5_GWAS).

### ASE-V association with Asthma and red blood distribution width

We tested the contribution of ASE associated SNPs to the two most significantly associated phenotypes-asthma and ‘red cell distribution width’ by assessing the ASE-variants and genes associated with these phenotypes (see Supplementary Table S6).

The strongest association between SNVs associated with a phenotype and ASE-V is for asthma (OR=1.50, p-value=1.27E-10). Positioning the significantly associated ASE-V on all chromosomes (Fig. 5B) shows that the entire signal is associated with the MHC locus. No other ASE-V outside of this locus meets the conservative GWAS genomic threshold (p-value=5.0E-08, Fig. 5C, dashed red line). A zoom-in to Chr6, ∼4M bp (29.5M to 33.5M) highlights a large collection of ASE-V variants that overlap class I and II of the MHC locus (Fig. 5C, bottom). The asthma ASE-V associated variants are shown in Supplemental Table S6_Asthma. The dominated signature at the MHC applies for all other listed ASE-associated AID including the unified phenotype of IBD/Crohn’s Disease, T1D, rheumatoid arthritis, and multiple sclerosis. However, the identity of the variants that specify each phenotype is mostly disease-specific (Supplemental Table S7).

Inspecting the associated ASE-V for ‘red cell distribution width’ shows that while a large fraction of the signal is also associated with the MHC loci (59 out 82 most significant associations with p-value <5E-08 (Supplementary Table S6_Red-blood), the rest of the ASE-V are distributed on several other chromosomes, with some ASE-V that are clustered (Fig. 5D, arrows). Among these variants, rs4246215 (chr11:61564298) is the most significant variant not included in the MHC locus (with p-value, 1.45E-57, Supplementary Table S6_Red-blood). We seek explanatory association of ‘red cell distribution width’ trait and this variant. According to results of the GWAS analysis for the 48 phenotypes from UKBB (Supplemental Table S5), rs4246215 is also strongly associated with platelets counts (p-value, 5.12E-47) and mean platelet volume (p-value, 1.11E-50) traits. Interestingly, GWAS results performed on an independent cohort connect this variant and its target gene FEN1 to a new function of the myeloid progenitor cell, in megakaryopoiesis and platelet formation (Gieger et al. 2011). Taking all together, we propose an explanation for the connection of blood cell phenotypes, being ASE and the function of FEN1.

### ASE genes enriched for processes related to viral infection

AlleleDB provides a list of 874 genes that are annotated as ASE genes (ASE-G) according to data combined from individual genomes, RNA-seq and Chip-seq (Chen et al. 2016). AlleleDB uses a strict statistical threshold for assigning genes as ASE. Fig. 6A shows the results for Gene Ontology (GO_biological processes) enrichment test for the ASE-G. The dominant enriched GO terms are associated with a response to interferon-gamma (GO:0060333, FDR q-value 5.48E-03, enrichment 10.66) and antigen processing and presenting of peptide antigen (GO:0048002, FDR q-value 9.79E-3, enrichment 9.48). A focused view on the interferon-gamma mediated signaling is shown in Fig. 6A.

**Figure 6.**
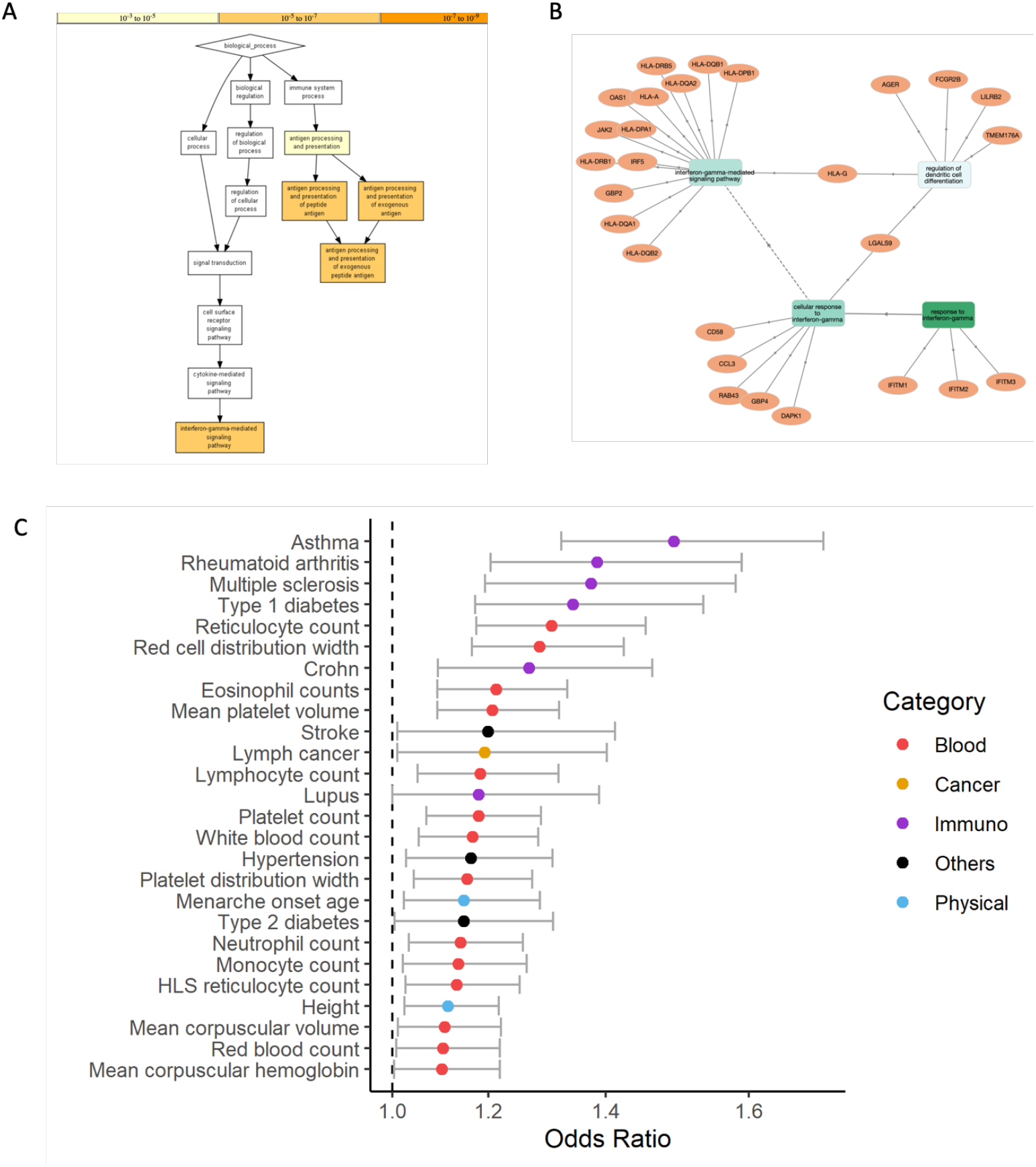
Enrichment of ASE genes for biological processes. **(A)** A graphical view for the results of an enrichment test for the 874 genes annotated as ASE-G. The test was performed by GOrilla enrichment tool (Eden et al. 2009). Interferon-gamma (GO:0060333) and antigen processing and presenting of peptide antigen (GO:0048002) are listed with high statistical confidence (see color codes for the significance of the enrichment by p-values range above the figure). **(B)** A network view for the GO terms and their associated genes (GOnet (Pomaznoy et al. 2018)). The enriched genes are shown with their gene symbols and the connection to the GO term for biological processes. **(C)** A list of traits and studies from the GWAS catalog that were significantly enriched by ASE (adjusted p-value<=0.1). All available GWAS studies and their associated genes were included in the test according to enrichment test of the GWAS catalog 2019 (Kuleshov et al. 2016). The traits are ranked by the log of the odds ratio (OR).

A network view of GO annotations for the interferon-gamma response (Fig. 6B) shows that a central gene hub includes many genes from the MHC locus (e.g. HLA-A, HLA-B). Additional genes are known to act in the viral activation (IRF-5), and the interferon response (e.g., GBP2, JAK2, OAS1). Specifically, OAS1 gene responds to interferon and consequently, influences viral RNA degradation and its replication cycle. Interestingly, polymorphisms in OAS1 have been associated with susceptibility to viral infection and T1D (Field et al. 2005). The expanded network includes other regulatory genes of the immune system. For example, LGALS9A which acts in modulating cell-cell and cell-matrix interactions. Altogether, the enrichment test for ASE-G shows evidence for the involvement of ASE-G in multiple aspects of viral infection cellular response.

### ASE genes signify a risk for a broad spectrum of immunological phenotypes by GWAS

Enrichment test for ASE-G versus the associated gene from the GWAS catalog was performed. The GWAS catalog is based on summary statistics for ∼72,000 variants and >5000 different GWAS reports that cover 1200 phenotypes. Fig. 6C ranks the significant traits from all available GWAS according to the calculated odds ratio (OR is in log scale). Supplemental Table S8 lists the enrichment and the statistically significant values for all results with adjusted p-value of <0.1.

A unified biological basis for most of these phenotypes is the strong immunological signature. For example, Neuromyelitis optica (including AQP4-IgG-positive) is an inflammatory disorder that is characterized by recurrent episodes of optic neuritis. While the involvement of specific cell types in activating these episodes remains unknown, the variability of the clinical manifestations according to the racial group, age of onset, the number of episodes (Palace et al. 2019) reflect phenotype diversity that might be governed by an individual ASE-V genomic makeup. Moreover, a recent GWAS that tested the intensity and pattern of mosquito bite revealed predisposition of alleles that are in association with T-cell regulation (Jones et al. 2017). Several of the listed significantly associated phenotypes are directly related to viral infection, including Hepatitis B virus (HBV) that affects the liver, Shingles that is caused by varicella-zoster virus (VZV), and Plantar warts that is caused by human papillomavirus (HPV). In summary, the listed ASE-genes associated phenotypes include mostly viral and bacterial infection, autoimmunity, antibody production and inflammation.

## Discussion

All cells in the human body share the same genome (exception are cells of the adaptive immune systems and germ cells) which is differentially expressed in tissues and cell types. In this study, we aim to use the collection of ASE-V from AlleleDB to reveal new insights on features that characterize the genomic occurrences of ASE-V. Assessing the extent of ASE across diploid organisms with current sequencing technology reveals that the unbalanced allele expression is a broader phenomenon than anticipated, and accounts for a substantial fraction of the transcriptome (Consortium 2015; Kita et al. 2017; Schmiedel et al. 2018). The unstable estimates for the extent of ASE from blood samples reflect the variability in cell composition and clonality of the immune cells in the blood (Schmiedel et al. 2018).

For this study, we have used the AlleleDB which reports on ∼10% on the expressed genes to be ASE genes (874 genes, 4% of all human coding genes). The data is based on ChIP-seq or RNA-seq associated with 382 individuals, and it is restricted to expression from LCL (Chen et al. 2016). Notably, in creating AllelDB (Chen et al. 2016) each sample was aligned to its personal reference individual haplotype eliminating many of the mapping pitfalls.

### Evolution selection constantis by ASE

Within AlleleDB ASE-V, when the dominating expressed allele was compared with the allele frequency reported in GenomAD (compiled from the 1000 Genome project and exomes sequencing of 60,000 healthy individuals), a strong trend toward expression from the major allele is observed (Fig. 4). This trend is not a mere reflection of the overall allelic expression pattern, as Acc-V that are not part of the ASE phenomenon show a minimal deviation from the exception. A similar trend towards the major allele applied to 90% of the instances in flycatchers (Wang et al. 2017). Whether ASE-V leans towards the major allele is a general trend in all organisms and tissues is yet to be determined.

The other feature for ASE-V concerns the increasing likelihood of ASE-V to dense polymorphic sites (Fig. 3). Both these features might indicate the potential of changes in the genome affecting the allelic expression choice (i.e. being expressed or not). We propose that ASE might be a mechanism to increase variability due to high polymorphism, while at the same time, maintaining stability by expressing the major allele. Variants of the major alleles in the population are the most likely ones to reflect the results of purifying selection. This, in turn, creates an allele-dependent fitness. Specifically, in different cell types, the relevant allele can be selected to support a desired phenotype (Schmiedel et al. 2018). At the individual level, ASE expression interacts with other genetic or environmental factors that may alter the allele expression choice thus affecting the outcome for complex phenotype and diseases (Buil et al. 2015; Edsgärd et al. 2016; Moyerbrailean et al. 2016)

A special case of evolution concerns the MHC and other immune related loci (Table 1). Specifically, the host-pathogen coevolution, maternal-fetal interaction (Hedrick 1998) are only some of the factors that shaped the human immune system (Borghans et al. 2004; Phillips et al. 2018). The system evolved to address conflicting objectives such as diversity, specificity, adaptability and self-tolerance. From a genomic perspective, the immune system is signified by high polymorphism (within MHC Class I and Class II and beyond (Norman et al. 2017)), gene clusters, and polymorphic frozen blocks (Gaudieri et al. 1997). The contribution of gene expression and specifically allele choice in cases of ASE in shaping the efficacy of the immune response, defining disease susceptibility and even in transplantation outcomes was only sporadically considered (see (Dawkins and Lloyd 2019; Petersdorf and O’HUigin 2019)).

### Diversity of phenotypes associated with ASE

Understanding the genetic basis for complex traits by GWAS is very challenging. Only rarely the associated variants are explanatory to the phenotype. The association between any specific allele and the phenotypic outcome dependents on numerous parameters (Boyle et al. 2017). We have tested the association of ASE-V to 48 major phenotypes including common diseases based on GWAS from the UKBB population (e.g. height, type 2 diabetes, Fig. 3). For many of these phenotypes we analyzed, 1000s of variants are expected to be associated with the trait, each with a minor effect size (Bomba et al. 2017). Our result show that ASE is beneficial for narrowing down the association when near-gene variants only are considered (see Methods). By focusing on ASE, we found that in contrast to many physical or metabolic phenotypes, the strongest ASE-V associated signal is to immune-related diseases and the properties of blood cells. For an inflammatory disease (asthma) and AID (e.g. T1D), the majority of the associated risk is confined to the MHC locus as reported before (Wandstrat and Wakeland 2001; Gutierrez-Arcelus et al. 2016). For the blood cell phenotypes, we observed that ASE-V is also dominated by the MHC locus along with additional genomic loci (Supplemental Table S7). The ASE associations to blood cells phenotypes cover properties of leukocytes, reticulocyte platelets, monocytes and more. In accord with of our results that identified ASE gene FEN1 as associated with blood cell phenotypes (rs4246215), it was shown in mice that mutations in this gene led to autoimmunity, chronic inflammation and cancers (Zheng et al. 2007). The shown examples and the shared ASE-V expose a genetic basis that unifies immune related phenotypes with the broad collection of blood cell phenotypes.

From all UKBB GWAS phenotype associations (48 total phenotypes), the association of ASE-V with asthma was maximal (OR=1.5). Asthma is an inflammatory disease that is associated with T helper (TH2), IgE production, and involvement of eosinophils (Holgate 2012). Despite a poor understanding of the genetic basis for asthma, overlap in the associated variants at MHC Class II was reported by other cohorts (Ramasamy et al. 2012). Moreover, a strong ASE signal is associated with major AID that affect a substantial fraction of the population (e.g. rheumatoid arthritis). One of the most studied AID is type 1 diabetes (T1D). The research of T1D is based on ∼40 large cohorts GWAS and discovered many risk SNVs and associated genes (Buniello et al. 2019). More than 50 genes were significantly associated with an increase or decrease the risk for T1D, with no proposed role for most of them, leaving T1D without a sufficient explanation to its etiology (Pociot and Lernmark 2016). Nevertheless, in many of these studies, MHC Class II locus and specific haplotypes (e.g., HLA-DR3-DQ2 and HLA-DR4-DQ8) were implicated as T1D genetic risk. We show that ASE-V and ASE-G are strongly associated with T1D but also with other major AID (e.g. rheumatoid arthritis, psoriasis, Crohn’s disease). We claim that a view through the lens of ASE-V might provide a new perspective on the clinical manifestation of AID and AID shared etiology.

The ASE-G that were identified in the enrichment tests (Figs 6A, 6B) include the building blocks of cell recognition by the immune cells (e.g. HLA, IG, Leukocyte IG receptors). Thus, phenotypes of autoimmunity, the propensity to viral and bacterial infection, and lymphoma are likely outcomes. Indeed, the broad range of traits that were significantly enriched by ASE-G cover different branches of the immune system. Most prominent traits (Fig. 6C) concerns the sensitivity of the immune system to fight viruses and bacteria. The enrichment for GO terms to interferon-gamma and antigen presentation on T-cells (Fig. 6A) provides a lead to the ASE involvement in the mechanism of cellular response to viral infection. In this view, the unexpected observation showing that different cell types of the immune system exhibit a unique and characteristic ASE profile is intriguing (Schmiedel et al. 2018), enabling a mechanism to maintain functional stability for the immune system. Personalized ASE profile might serve as a sensor for assessment of susceptibility to viral and bacterial infection, allergy, autoimmunity, inflammatory condition and even immunization efficacy. In this view, ASE-V for HLA and IG genes were suggested in identifying high-risk individuals for planning targeted immunotherapy (Liu et al. 2019). Along this line, chronic activation of the immune system (e.g. by viral or parasite infection, allergies) can lead to alteration in the ASE profile of the exposed cells, as illustrated by in-vitro stimulation of cells (Edsgärd et al. 2016). In summary, our results connecting the ASE with human phenotypes suggest that unbalanced expression of genes by ASE contributes to individual genetic variability toward complex traits and diseases, with major implications on immune-related conditions.

## Methods

### Database

The core data source of this research is AlleleDB available from the Gerstein lab (Chen et al. 2016). The database consists of heterozygous SNVs (hSNVs) and quantified data on allelic expression. The data combine large-scale RNA-seq and Chip-seq experiments that were performed on lymphoblastoid cell lines (LCL). The analyzed 382 individuals are accompanied by complete genomic information as being part of the 1000 Genomes Project. Each transcriptomic analysis has been aligned against its personal genome constructed from the autosomal chromosomes of GRCh37 (hg19) genome. Various additional filtration steps and technical protocol for harmonization are described in (Chen et al. 2016). Briefly, the dataset consider for each hSNVs from all individuals (n=5,284,835) the expression level from each of the alleles, along with binomial and beta-binomial statistics. Based on the quality levels and the deviation from the beta-binomial model, each hSNV is labeled as an ASE variant (ASE-V) or not. We adhered to the AlleleDB labels of Accessible SNV and ASE SNVs (abbreviated Acc-V and ASE-V, respectively) (Chen et al. 2016). The SNVs that are labeled ASE (a subset of Acc-V) are organized in a separate table (n=363,993). Evidence for Acc and ASE variants are based on an average of 8.76 and 5.22 individuals, respectively. For our analysis, we eliminated variants that are supported by only a single individual. For all further analysis, a random selection of an SNV representative is used per site. Remaining overall 287,449 Acc-V and out of them, 29,346 are ASE-V.

### Chromosomal analysis using windows

Each autosomal chromosome length (N) was extracted from the official site of the Genome reference of NCBI for GRCh37 (hg19). The chromosomes were segmented to windows to study them at different resolutions. Windows used are with length (L): 100 bp, 1000 bp, 10,000 bp and 100,000 bp. For each window, the number of Acc-V and ASE-V were counted in an overlapping window with L/2 between consecutive windows. For each window length, a cumulative distribution along the chromosome for ASE-V and Acc-V occurrences is calculated. Kolmogorov-Smirnoff of two sample test (KS test) is applied for the cumulative distributions for each chromosome separately. For 100k bp segments, a value of the fraction of the ASE-V out of Acc-V is calculated as:

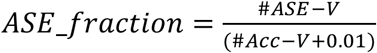

### Spacing estimates

For the spacing distribution test, we measure the minimal distances in bp between any consecutives Acc-V compared to the expected distances. We applied the test for Chr1 (total length of 250M bp). The effective length of Chr1 transcriptome (T) is estimated from the number of 10k bp windows with Acc-V for Chr1 as a representative. The fraction of positive occurrences of Acc-V segments (11.2%) was used to estimate Chr1 effective length of transcription (T, 28M bp). All together we measured from the uniquely compressed Acc-V set (as reported in (Chen et al. 2016)) N distances (N=5835 distances for Chr1, Supplemental Table S3). The expected Acc density (d) is defined by d=N/T. We calculated the expectation of observing a distance that is bounded by any length (L). The geometric model is expected to follow a value of *N*(1 − (1 − *d*)^*L*^). The calculated values by the model were compared to the actually observed distances.

### ASE occurrence dependency

We quantify the probability of Acc-V to be labeled as ASE-V given the inherent density of the Acc-V (i.e. being expressed in the sample). Altogether, for all autosomal chromosomes, 10.21% of the Acc-V are annotated as ASE-V (29,346 out of 287,453 unique sites). The tendency of ASE clustering is calculated for a predefined length (e.g., L=100 bp). We recorded the observed fraction of ASE-V according to the predefined number of Acc-V. For all windows of 100 bp with exactly k Acc-V within (k is 1,2,3..), we asked for how many of them with an instance of ASE (>=1) exists. For a theoretical calculation under the assumption of independent occurrences, we calculated the expected value to be: 1 − (1 − *p*^*k*^), where p is the fraction of ASE-V/Acc-V in the genome (0.1021), and k is the selected number of Acc appearance. We repeated the same analysis for windows of size 1000 bp.

### ASE preference

For each Acc-V, we checked in AlleleDB, which allele (the Reference or the Alternative) had a higher expression according to the reads counted for each allele. The allele tendency was recorded for each of the Acc-V as being 1 for the Alt allele, or 0 for the Ref allele. Additionally, each SNV has been associated with its alternative allele frequency (AAF) according to the data collected from Genome Aggregation Database (GnomAD) (Karczewski et al. 2019) (as reported in gnomad.exomes.r2.0.2 VCF file). The SNVs were binned according to their AAF in windows of with an interval of 0.05 so that each bin corresponds to a range of AAF values (e.g. 0-0.05). For each of these bins, the mean for being transcribed from the Alt allele was calculated both for the ASE-V and the complementary set of Acc-V (all Acc-V excluding ASE-V). Logistic regression was used to determine the correlation between a variant which is being transcribed from the Alt allele to the actual AAF value as reported by healthy human population according to GnomAD.

### UK Biobank GWAS analysis

The GWAS was performed on a cohort derived from the UK Biobank (UKBB) dataset (Sudlow et al. 2015; Bycroft et al. 2017). From the entire UKBB cohort of 502,539 samples with genotypes, information of ethnicity and gender was taken. To avoid biases due to family relationship, we removed 75,853 samples to keep only one representative of each kinship group of related individuals. Additional 312 samples had mismatching sex (between the self-reported and the genetics-derived) and 726 samples with only a partial genotyping. The filtered collection includes 332,709 samples. The list of phenotypes on which we performed GWAS is available in Supplementary Table S5. The table specifies how each phenotype was defined (based on a UKBB field or an ICD-10 code). The set of all ICD-10 codes associated with a sample were derived from the following UKBB fields: 41202, 41204, 40006, 40001, 40002, 41201.

The following covariates were included in the analysis: sex (binary), year of birth (numeric), the 40 principal components of the genetic data as provided by the UKBB (numeric), the genotyping batch (one-hot-encoding with 105 categories) and the assessment centers associated with each sample (binary, with 25 categories). Altogether, 173 covariates (including the intercept) were included in the analysis. For each phenotype, we tested all 804,069 genetic markers genotyped by the UK Biobank Axiom Array, variants provided by the UKBB (Bycroft et al. 2017). Continuous phenotypes were analyzed by a multivariate linear regression model, and binary phenotypes by a logistic regression model (using the Statsmodels Python package (Seabold and Perktold 2010)).

For each of the listed phenotypes, we performed a Fisher exact test for enrichment of ASE-V in the SNVs that are associated with the phenotype at a significant level of p-value <0.05. Overall, the overlap of the two experiments SNVs included 34,055 Acc-V and 2,631 SNVs that are ASE-V. OR and p-values were calculated for each of the listed 48 phenotypes.

### Enrichments tests of ASE genes

A complete list of genes labelled as ASE genes was downloaded from the compiled list of all expressed genes (Chen et al. 2016). The list consists of 20,142 genes. Out of them, 874 are considered ASE genes. For the ASE genes list we tested by GOrilla enrichment tool (Eden et al. 2009) for GO_Biological Process (GO_BP) terms. GOnet was used to provide a network view by graph representation for the genes enriched by the GO_BP annotations (Pomaznoy et al. 2018). Enrichment for the GWAS catalog 2019 (Buniello et al. 2019) was done according to the analysis of Enrichr for the list of 874 ASE genes according to default human parameters (Kuleshov et al. 2016). The output includes the number of genes that are associated with ASE and a phenotype out of the total genes associated with this phenotype. We used these values along with the overall number of expressed genes in AlleleDB, to calculate the Fisher Exact test, OR and p-values for the association of each of the phenotypes with ASE.

## Acknowledgment

Special regards to Jieming Chen from Gerstein’s lab for the kind answers to numerous technical questions. We thank Nati Linial (Hebrew University) for insightful comments and for support in the probabilistic analysis. We thank Liran Carmel and Sagiv Shifman (Hebrew University) for useful comments. We thank all members of the Linial lab for helpful discussions throughout the project. Special thanks for Nadav Brandes that provided the results of an in-house GWAS performed on the UK Biobank (under the UKBB application ID 26664), and for his professional support on data analysis. Special thanks for the CS system for computational infrastructure and services. KWK is a recipient of the Ariane de Rothschild Scholarship.

## Data access

All the analyses in this study were based on published data sets. A tab-delimited spreadsheet containing all supporting filtered sets, GWAS inhouse results and other related data generated in this study is provided in Supplemental Tables 1-8. The full results of the GWAS analysis on UKBB, including detailed quality-control reports, will soon be published elsewhere. We will provide the summary statistic of the results upon demand (under UKBB application ID 26664).

## Contributions

KWK and ML conceived the method, developed the analysis scheme, and wrote the manuscript. KWK formally led the analyses on AlleleDB and conducted the statistical tests. Both authors read and approved the final manuscript.

